# SEXUAL DIMORPHISM IN FIN SIZE AND SHAPE IN BLUEFIN KILLIFISH

**DOI:** 10.1101/2025.11.18.689072

**Authors:** Kasey Brockelsby, Elijah J. Davis, Olivia A. Roden, Valerie Shamsyna, Rebecca C. Fuller

## Abstract

Sexual dimorphism provides insight into how trait optima differ between males and females, despite their shared genome. Measuring sexual dimorphism can help identify which traits have been shaped by sexual selection. While fish morphology has been widely described, fewer studies have quantified sexual dimorphism across all fin types—pectoral, pelvic, dorsal, anal, and caudal. Fins are often overlooked due to their small size, tendency to fold against the body, and poor preservation post-collection. In this study, we quantified sexual dimorphism in fin size and shape across all fin types in the bluefin killifish, *Lucania goodei*. We found striking sexual dimorphism in the dorsal and anal fins, particularly in area, ray length, and base length. In contrast, the pelvic, pectoral, and caudal fins showed moderate, but detectable, levels of dimorphism. In both dorsal and anal fins, males exhibited elongation of the posterior region relative to females. Dorsal and anal fin traits (i.e., area, ray length, base length) were also strongly correlated within both sexes, such that individuals with larger than average dorsal fins also have larger than average anal fins. This correlation is present for both males and females, suggesting shared developmental pathways, pleiotropy, or correlated selection between dorsal and anal fins. Overall, our results indicate that the dorsal and anal fins are key targets of sexual selection in males, likely reflecting their roles in courtship, competition, and/or external fertilization.

## INTRODUCTION

Sexual dimorphism is common across fishes (Breder and Rosen, 1966; Andersson, 1994). Males and females often differ in coloration, presence/absence of breeding tubercles, body size/shape, vocalizations, and behavior. Fewer studies have focused on sexual dimorphism in the size and shape of fins, although noticeable exceptions exist in some groups (Im, et al., 2016; Goldberg, et al., 2019; Sowersby, et al., 2022). Fins are intriguing because they are multifunctional. They affect multiple aspects of swimming, including stabilization in the water column, forward thrust, and turning (Standen and Lauder, 2005; Flammang and Lauder, 2016). Fins are also utilized in mating, including the courtship of females, displays towards competing males, and the movement and coordination of mating during pair spawning (Breder and Rosen, 1966; Basolo, 1990; Rosenthal and Evans, 1998; Kozak, et al., 2008). Some species even use their fins to ‘clasp’ females during spawning (Able and Hata, 1984; Sabaj, et al., 2000; Hayakawa and Kobayashi, 2010).

A sizable literature examines the evolution of fish body morphology, fin placement, and length of spiny rays as a function of ecology (Langerhans, 2008; Chang and Alfaro, 2016; Price, et al., 2019). However, less attention has been given to the fins, particularly to the soft ray fins (Standen and Lauder, 2005). Soft-rayed fins often lie flat against the body when fish are removed from water. Additionally, geometric morphometrics techniques are more challenging to apply to soft ray fins since they have fewer non-flexible landmarks. Despite these challenges, sexual dimorphism in fin size and shape is documented, including the extreme dimorphisms among livebearing fishes, with the enlarged dorsal fin of sailfin mollies (*Poecilia reticulata*) and the elongated caudal fin found in sword tail fishes (*Xiphophorus*) (Basolo, 1990; Rosenthal and Evans, 1998; Rosenthal, et al., 2001; Kozak, et al., 2008). Other members of the Poeciliidae are also known for having dimorphic dorsal fins (Goldberg, et al., 2019). However, sexual dimorphism in fin traits may be common across the order Cyprinodontiformes, to which poeciliids belong. Recently, Davis et al. (2025) measured sexual dimorphism across 20 species of North American killifish (Fundulidae) and found pronounced dimorphism in dorsal and anal fin traits, but modest dimorphism in caudal fin traits (see also Welsh, et al., 2013; Welsh and Fuller, 2015). Pronounced sexual dimorphism in dorsal and anal fin traits has also been found in other families of killifish (Sowersby, et al., 2022; Mainero, et al., 2023). A closely related order (Beloniformes) contains the medaka, which also possesses pronounced dimorphism in dorsal and anal fins in some populations (Sumarto, et al., 2020; Downer-Bartholomew and Rodd, 2025).

Numerous studies have examined sexual dimorphism in a subset of the fin traits. Sticklebacks and rock fish have been well-studied with regard to sexually dimorphic pectoral fins (Echeverria, 1986; Bronseth and Folstad, 1997; Bakker and Mundwiler, 1999). Similarly, some cods, cichlids, and whitefish have sexually dimorphic pelvic fins (Brichard, 1989; Karino, 1997; Casselman and Schulte-Hostedde, 2004; Skjæraasen, et al., 2006). Mainero et al. (2023) examined the length of fins in the Arabian killifish, *Aphaniops stoliczkanus,* and found sexual dimorphism in anal, dorsal, pectoral, and pelvic fin lengths and base lengths when scaled to standard length. Anal and dorsal fins had particularly high levels of sexual dimorphism in comparison to the other fins.

Fins generally provide an excellent tapestry for investigating classic questions regarding natural and sexual selection and the nature of genetic constraints. As multifunctional traits, they impact swimming and maneuvering and are often used in signaling and mating. As morphological traits, they also provide a compelling system to pose basic questions about trait correlations and trait variability (Falconer, 1996). To what extent is fin growth and size correlated across multiple fin types? What are the overall levels of trait variability? Are sexually selected traits more or less variable than other traits? Do levels of variability differ between males and females? These questions are central to understanding the evolutionary forces shaping morphological traits. Some theories predict that condition-dependent traits, which are often exaggerated in males, should exhibit high variability (Rowe and Houle, 1996; Houle and Kondrashov, 2002; Zajitschek, et al., 2020). Conversely, traits under strong directional or stabilizing selection are expected to have low levels of genetic variation. By comparing variability across traits and between sexes, we can assess whether patterns of variation align with predictions based on condition-dependent sexual signaling and general selective pressures (Kodric-Brown and Hohmann, 1990).

This study sought to estimate the levels of sexual dimorphism across all fin types (dorsal, anal, caudal, pectoral, and pelvic) and to determine the relative levels of variability and covariation. To answer these questions, we focused on the bluefin killifish, *Lucania goodei*, which are known for displaying high levels of sexual dichromatism in their fins; males have brightly colored dorsal and anal fins (Foster, 1967; Fuller, 2002; Fuller, et al., 2022) that are signals of condition and dominance (Johnson and Fuller, 2015). The pelvic fins and the caudal fin base are also brightly colored in males. By contrast, female bluefin killifish are drab, with minimal fin color. The pectoral fins of both species exhibit no color at all in either sex.

In this study, we asked the following: Which fins show the highest levels of sexual dimorphism in size and/or shape? Do fins with sexual dichromatism (i.e., male nuptial colors) have higher levels of sexual dimorphism in size and shape? Do males have higher levels of trait variability than females in fins with high levels of dimorphism? Are there tight correlations between trait values that suggest pleiotropy and/or a shared developmental pathway? To answer these questions, we took high-resolution pictures of male and female bluefin killifish (*Lucania goodei*) and measured multiple aspects of the dorsal, anal, caudal, pelvic, and pectoral fins. We found that dorsal and anal fins are particularly dimorphic and covary with one another, suggesting that males with large dorsal fins also have large anal fins, even after controlling for overall size. We also found high correlations between dorsal and anal fin traits in both males and females, suggesting the presence of pleiotropy, shared developmental pathways, and/or correlated selection.

## Methods

Our goal was to measure sexual dimorphism in fin size and shape in the bluefin killifish, *Lucania goodei,* to determine whether sexually dimorphic traits had higher (or lower) levels of variability, and to determine whether there were strong correlations between different fin elements. Fish were collected from the wild from two different populations in Florida: Rainbow River (clear, spring-fed river, Marion County) and Everglades 26 Mile Bend (swamp, Broward County) populations, in May 2021. The fish were returned to the laboratory and maintained in a greenhouse until Spring 2022. We measured fin size, shape, and standard length for approximately 10 males and 10 females from each population. Individuals were sexed based on diagnostic sexually dimorphic characteristics, namely the presence of color and black melanin spots and/or borders on the dorsal and anal fins of males. Fish were euthanized in a buffered solution of MS-222 (tricaine mesylate) so that their fins could be manipulated more easily and then photographed.

Fish were photographed in a petri dish with a small amount of water on a plain grey stage, using a Nikon D5600 camera with an AF-S Micro Nikkor 105mm lens. To minimize glare and ensure overall quality, two lamps were angled to illuminate the stage with diffuse light. Finally, a small ruler was placed on the stage underneath the petri dish to provide a scale, and male fish were also photographed with a color standard in the frame. At least four photographs were taken of every fish: one photo capturing the full body of the fish, then one photo focused on each unpaired fin (dorsal, anal, and caudal). Once these photos had been taken, the pectoral and pelvic fins of the fish were cut off with small dissecting scissors and placed on a microscope slide for further photography. Microscope slides were placed on the grey stage under the same lighting conditions with a small ruler for scale. We took photos focused on the pelvic and pectoral fins.

Measurements were obtained using ImageJ ver. 1.53k. We measured standard length (tip of the snout to the base of the caudal fin), length of the base of each fin (insertion of the first ray to the insertion of the last ray), lengths of individual fin rays within each fin, and area of each fin. The number of rays in each fin was also counted. Figure 1 shows an example of linear measurements taken on a specimen. Caudal fin ray counts and lengths were measured following the protocols described by Armbruster (2012). Specifically, we measured the median caudal ray and all branched caudal rays plus one unbranched ray adjacent to the branched rays on both the dorsal and ventral sides. These measurements were performed twice by separate people. We measured fin area, fin base length, number of fin rays, and fin ray lengths for pectoral and pelvic fins using the images of fins removed from the body.

**Figure 1.**
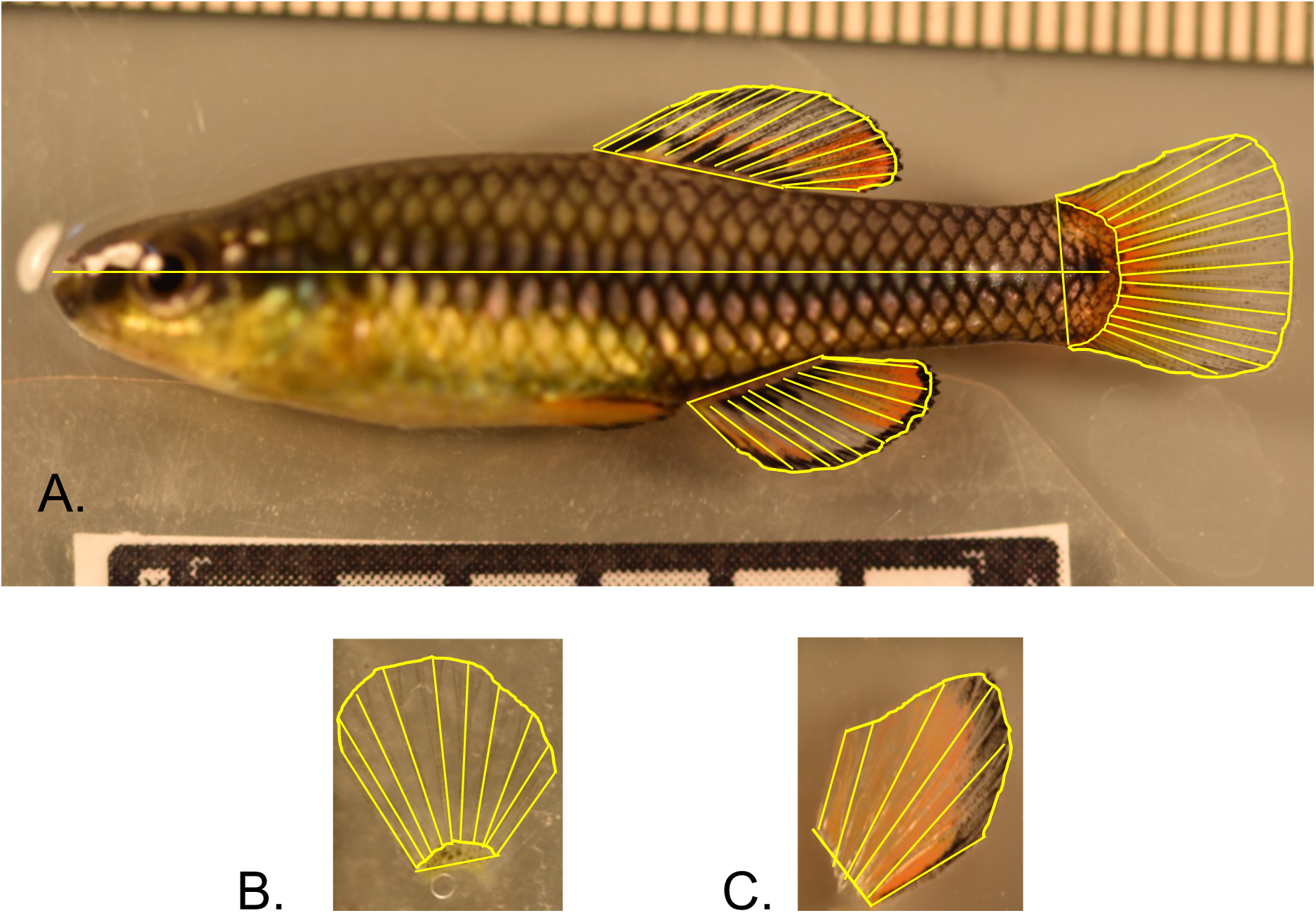
Male *L. goodei* with measurements shown for standard length and dorsal, anal, and caudal fins traits (fin base length, area, ray number, and ray length); B-C) pelvic and pectoral fin traits (base length, area, fin ray number, fin ray lengths). Photo credit Kasey Brockelsby.

### Data Analysis

We calculated the repeatability of measurements taken by multiple investigators and multiple photos of the same specimen. Dorsal and anal fin traits were measured by two separate people using two separate photos (i.e., four total measurements per trait per individual). All other traits were measured using a single photo measured by two people (i.e., two total measurements per trait per individual). Repeatability was calculated using the ‘lmer’ function from the ‘lmer4’ library in R, where we treated each individual fish as a random factor.

The overarching goals of this project were to determine (1) which fins (i.e., dorsal, anal, caudal, pectoral, or pelvic) and fin traits (i.e., base length, ray length, area) show the highest levels of sexual dimorphism, (2) whether some fins are particularly variable and whether this varies as a function of sex, and (3) whether there are strong correlations across these traits independent of size. We considered individuals as the unit of replication. Multiple measurements of the same trait on the same individual were averaged to produce a single value. For each fin, we also calculated the mean ray length for all rays. Fin area, fin ray lengths, and base lengths were all positively correlated with standard length. To examine fin size and shape independent of overall size, we calculated the residuals from linear regressions of fin areas, base lengths, and ray lengths against mean standard length. We then used a linear model to test for differences in residual trait variation due to sex, source population, and an interaction between the two. Only one trait varied significantly as a function of the interaction between sex and population, and this test did not remain statistically significant after a sequential bonferonni correction. Hence, we ran a simple additive model that considered the effects of sex and population, but not their interactions.

Sexual dimorphism was measured as the effect size due to sex. The effect was measured with the ‘eff_size’ function from the ‘emmeans’ library in R. This is the difference in means between the two sexes divided by the pooled standard deviation. We also calculated the 95% confidence limits to ask whether sexual dimorphism differed from zero and whether it differed from other traits.

To visualize sex differences in fin size and shape, we calculated the least-square means and standard errors for each ray length and the base length for all the fins from separate models that considered the effects of standard length and sex on trait value. To visualize the fins, the first fin ray was located at the origin (0), the last ray was located at the end of the sex-specific fin base, and the intermediate rays were evenly spaced between the first and last rays. This allowed us to visualize sex-differences in base length, fin ray lengths, and shape.

To determine whether the two sexes differed in the levels of variability for a given trait, we estimated the sex-specific absolute means and coefficients of variation on the raw data as well as the size-corrected coefficients of variation. For the size-corrected coefficients of variation, we calculated the standard deviation of the residuals from a linear regression of trait value on standard length for both males and females and then divided the standard deviations by the absolute means.

We also investigated the correlations among size-corrected fin traits to determine whether there were particularly strong correlations among some fins and whether this pattern differed between males and females. To do this, we calculated the residuals from separate regressions of standard length on trait values for each sex and each trait. We then examined the correlations among fin area, fin ray lengths, and fin base lengths separately for each sex. For each fin trait (i.e., area, ray length, base length) and each sex, we performed a sequential Bonferonni correction.

All analyses were performed in R version 4.5.0. Raw data and R scripts can be found at http://datadryad.org/share/aI4FXI2axjfyHUiU-NmdJQId_o_NvTcfyPuRFxWx2C0.

## Results

Repeatability was generally high (average repeatability = 0.80, Table 1). Most traits were positively correlated with standard length, with an average correlation of 0.54 between standard length and continuous fin traits (i.e., fin area, fin ray lengths, and fin base length, Table 2). Ray counts were not positively correlated with standard length (*p* > 0.27), with the exception of dorsal ray counts (*R_37_* = 0.33, *p* = 0.034). Analyses of variance indicated that ray counts did not vary as a function of sex or population (*p* > 0.09 in all tests), regardless of whether standard length was taken into account. Finally, standard length varied as a function of population of origin (Figure 2, *F_1,35_* = 34.08, *p* = 1.263e-06), but there were no statistically significant effects of sex (*F_1,35_* = 0.13, *p* = 0.7236) or the interaction between sex and source population (*F_1,35_* = 0.31, *p* = 0.5830). For the remaining analyses, we consider size-adjusted trait values as the residual values of traits regressed on standard length.

**Figure 2.**
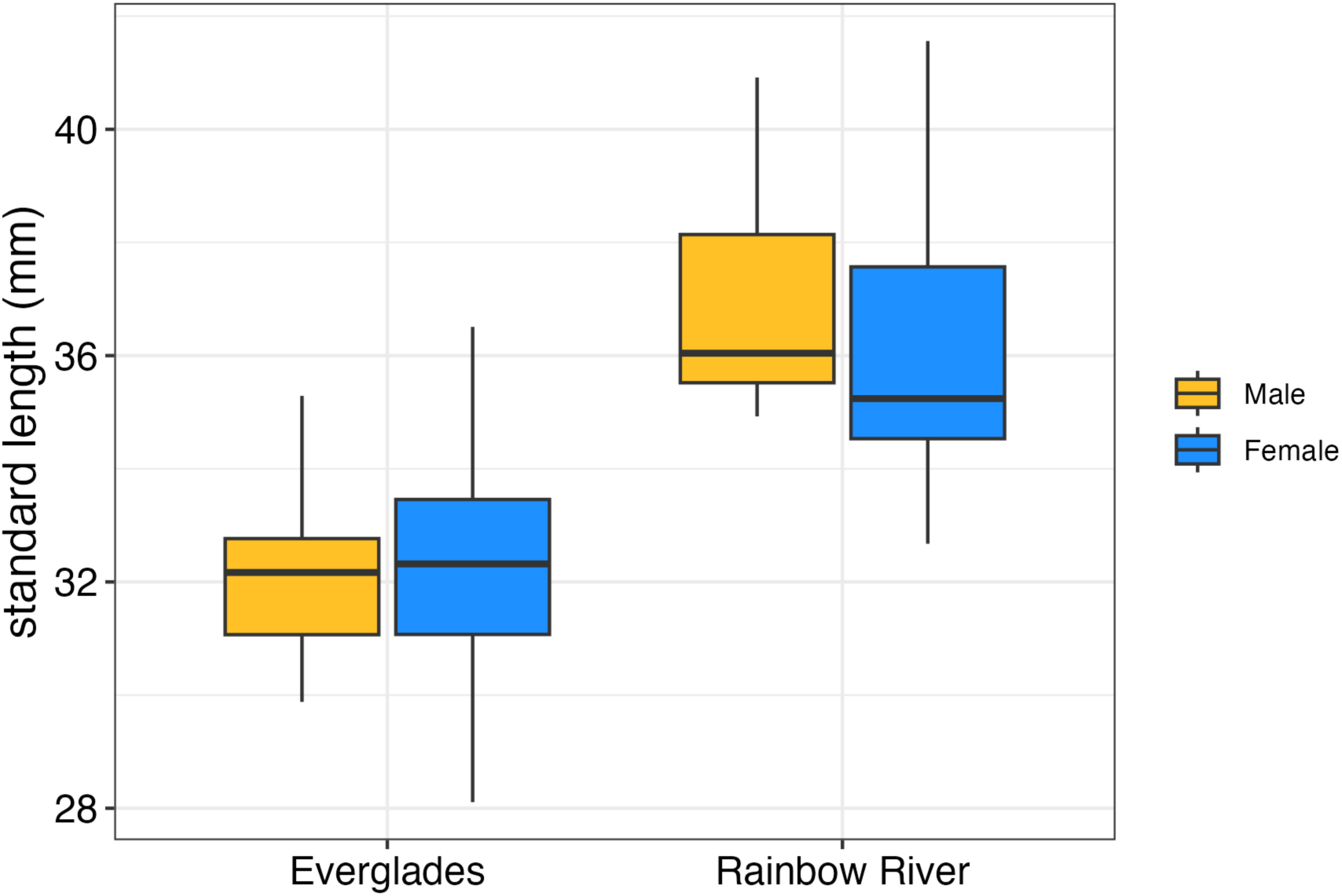
Standard length as a function of sex and population.

**Table 1.** Repeatability across traits. Repeatability is listed for traits where the sample size across individuals was greater than 20.

| **Traits** | **Repeatability** | **Total**<br>**Individuals** | **Total**<br>**Images** |
| --- | --- | --- | --- |
| Standard Length | 0.96 | 39 | 78 |
| Dorsal Rays # | 0.84 | 39 | 154 |
| Dorsal Area | 0.97 | 39 | 154 |
| Dorsal Base Length | 0.74 | 39 | 154 |
| Dorsal 1 Ray Length | 0.80 | 39 | 154 |
| Dorsal 2 Ray Length | 0.77 | 39 | 154 |
| Dorsal 3 Ray Length | 0.68 | 39 | 154 |
| Dorsal 4 Ray Length | 0.86 | 39 | 154 |
| Dorsal 5 Ray Length | 0.90 | 39 | 154 |
| Dorsal 6 Ray Length | 0.89 | 39 | 154 |
| Dorsal 7 Ray Length | 0.93 | 39 | 154 |
| Dorsal 8 Ray Length | 0.93 | 39 | 154 |
| Dorsal 9 Ray Length | 0.94 | 39 | 154 |
| Dorsal 10 Ray Length | 0.94 | 38 | 145 |
| Dorsal 11 Ray Length | 0.95 | 27 | 99 |
| Anal Ray # | 0.78 | 39 | 153 |
| Anal Area | 0.95 | 39 | 153 |
| Anal Base Length | 0.94 | 39 | 153 |
| Anal 1 Ray Length | 0.70 | 39 | 152 |
| Anal 2 Ray Length | 0.61 | 39 | 152 |
| Anal 3 Ray Length | 0.66 | 39 | 152 |
| Anal 4 Ray Length | 0.78 | 39 | 152 |
| Anal 5 Ray Length | 0.82 | 39 | 152 |
| Anal 6 Ray Length | 0.87 | 39 | 152 |
| Anal 7 Ray Length | 0.89 | 39 | 152 |
| Anal 8 Ray Length | 0.90 | 39 | 151 |
| Anal 9 Ray Length | 0.88 | 39 | 151 |
| Anal 10 Ray Length | 0.94 | 34 | 123 |
| Caudal Ray Number | 0.89 | 39 | 79 |
| Caudal Area | 0.96 | 39 | 146 |
| Caudal Base Length | 0.88 | 39 | 144 |
| Median Caudal Ray Length | 0.94 | 39 | 79 |
| Caudal Dorsal 1 Ray Length | 0.81 | 39 | 79 |
| Caudal Dorsal 2 Ray Length | 0.90 | 39 | 79 |
| Caudal Dorsal 3 Ray Length | 0.89 | 39 | 79 |
| Caudal Dorsal 4 Ray Length | 0.92 | 39 | 79 |
| Caudal Dorsal 5 Ray Length | 0.86 | 39 | 79 |
| Caudal Dorsal 6 Ray Length | 0.73 | 39 | 78 |
| Caudal Dorsal 7 Ray Length | 0.65 | 22 | 45 |
| Caudal Ventral 1 Ray Length | 0.73 | 39 | 79 |
| Caudal Ventral 2 Ray Length | 0.90 | 39 | 79 |
| Caudal Ventral 3 Ray Length | 0.90 | 39 | 79 |
| Caudal Ventral 4 Ray Length | 0.89 | 39 | 79 |
| Caudal Ventral 5 Ray Length | 0.86 | 39 | 79 |
| Caudal Ventral 6 Ray Length | 0.75 | 39 | 78 |
| Pelvic Rays # | 0.61 | 38 | 75 |
| Pelvic Area | 0.95 | 39 | 78 |
| Pelvic Base Length | 0.57 | 39 | 78 |
| Pelvic 1 Ray Length | 0.75 | 39 | 78 |
| Pelvic 2 Ray Length | 0.72 | 39 | 78 |
| Pelvic 3 Ray Length | 0.82 | 39 | 78 |
| Pelvic 4 Ray Length | 0.89 | 39 | 78 |
| Pelvic 5 Ray Length | 0.78 | 39 | 77 |
| Pelvic 6 Ray Length | 0.81 | 34 | 62 |
| Pectoral Rays # | 0.63 | 37 | 73 |
| Pectoral Area | 0.94 | 37 | 73 |
| Pectoral Base Length | 0.77 | 37 | 73 |
| Pectoral 1 Ray Length | 0.55 | 37 | 73 |
| Pectoral 2 Ray Length | 0.46 | 37 | 73 |
| Pectoral 3 Ray Length | 0.53 | 37 | 73 |
| Pectoral 4 Ray Length | 0.61 | 37 | 73 |
| Pectoral 5 Ray Length | 0.71 | 37 | 73 |
| Pectoral 6 Ray Length | 0.77 | 37 | 73 |
| Pectoral 7 Ray Length | 0.70 | 36 | 72 |
| Pectoral 8 Ray Length | 0.66 | 35 | 70 |
| Pectoral 9 Ray Length | 0.66 | 33 | 64 |
| Pectoral 10 Ray Length | 0.45 | 24 | 43 |

**Table 2.** Correlation coefficients between standard length and fin traits.

| **Trait** | **R** | **DF** | **P** |
| --- | --- | --- | --- |
| Dorsal Ray# | 0.33 | 37 | 0.0395 |
| Dorsal Area | 0.43 | 37 | 0.0065 |
| Dorsal Base Length | 0.59 | 37 | 7.62E-05 |
| Mean Dorsal Fin Ray Length | 0.51 | 37 | 0.0008 |
| Dorsal Ray 1 Length | 0.31 | 37 | 0.0566 |
| Dorsal Ray 2 Length | 0.51 | 37 | 0.0010 |
| Dorsal Ray 3 Length | 0.59 | 37 | 6.80E-05 |
| Dorsal Ray 4 Length | 0.57 | 37 | 0.0002 |
| Dorsal Ray 5 Length | 0.59 | 37 | 7.96E-05 |
| Dorsal Ray 6 Length | 0.54 | 37 | 0.0003 |
| Dorsal Ray 7 Length | 0.50 | 37 | 0.0012 |
| Dorsal Ray 8 Length | 0.46 | 37 | 0.0030 |
| Dorsal Ray 9 Length | 0.50 | 37 | 0.0011 |
| Dorsal Ray 10 Length | 0.54 | 36 | 0.0004 |
| Dorsal Ray 11 Length | 0.55 | 25 | 0.0028 |
| Dorsal Ray 12 Length | 0.44 | 2 | 0.5556 |
| Anal Ray# | 0.16 | 37 | 0.3344 |
| Anal Area | 0.50 | 37 | 0.0011 |
| Anal Base Length | 0.62 | 37 | 2.47E-05 |
| Mean Anal Fin Ray Length | 0.49 | 37 | 0.0016 |
| Anal Ray 1 Length | 0.16 | 37 | 0.3237 |
| Anal Ray 2 Length | 0.36 | 37 | 0.0245 |
| Anal Ray 3 Length | 0.43 | 37 | 0.0065 |
| Anal Ray 4 Length | 0.47 | 37 | 0.0025 |
| Anal Ray 5 Length | 0.47 | 37 | 0.0028 |
| Anal Ray 6 Length | 0.43 | 37 | 0.0060 |
| Anal Ray 7 Length | 0.44 | 37 | 0.0054 |
| Anal Ray 8 Length | 0.46 | 37 | 0.0030 |
| Anal Ray 9 Length | 0.46 | 37 | 0.0033 |
| Anal Ray 10 Length | 0.44 | 32 | 0.0084 |
| Anal Ray 11 Length | 0.37 | 15 | 0.1425 |
| Caudal Ray# | 0.05 | 37 | 0.7754 |
| Caudal Area | 0.65 | 37 | 8.68E-06 |
| Caudal Base Length | 0.84 | 37 | 1.94E-11 |
| Mean Caudal Fin Ray Length | 0.84 | 37 | 2.31E-11 |
| Median Caudal Ray Length | 0.84 | 37 | 1.98E-11 |
| Caudal Dorsal 1 Ray Length | 0.80 | 37 | 1.40E-09 |
| Caudal Dorsal 2 Ray Length | 0.85 | 37 | 1.24E-11 |
| Caudal Dorsal 3 Ray Length | 0.82 | 37 | 1.24E-10 |
| Caudal Dorsal 4 Ray Length | 0.78 | 37 | 3.80E-09 |
| Caudal Dorsal 5 Ray Length | 0.81 | 37 | 6.00E-10 |
| Caudal Dorsal 6 Ray Length | 0.71 | 37 | 3.74E-07 |
| Caudal Dorsal 7 Ray Length | 0.83 | 20 | 2.14E-06 |
| Caudal Ventral 1 Ray Length | 0.80 | 37 | 1.49E-09 |
| Caudal Ventral 2 Ray Length | 0.84 | 37 | 2.09E-11 |
| Caudal Ventral 3 Ray Length | 0.84 | 37 | 1.70E-11 |
| Caudal Ventral 4 Ray Length | 0.81 | 37 | 4.69E-10 |
| Caudal Ventral 5 Ray Length | 0.74 | 37 | 5.41E-08 |
| Caudal Ventral 6 Ray Length | 0.61 | 37 | 3.89E-05 |
| Caudal Ventral 7 Ray Length | -0.64 | 2 | 0.3646 |
| Pelvic Ray# | 0.18 | 36 | 0.2709 |
| Pelvic Area | 0.48 | 37 | 0.0018 |
| Pelvic Base Length | 0.33 | 37 | 0.0418 |
| Mean Pelvic Fin Ray Length | 0.48 | 37 | 0.0020 |
| Pelvic Ray 1 Length | 0.36 | 37 | 0.0241 |
| Pelvic Ray 2 Length | 0.50 | 37 | 0.0011 |
| Pelvic Ray 3 Length | 0.47 | 37 | 0.0023 |
| Pelvic Ray 4 Length | 0.42 | 37 | 0.0083 |
| Pelvic Ray 5 Length | 0.38 | 37 | 0.0171 |
| Pelvic Ray 6 Length | 0.27 | 32 | 0.1185 |
| Pectoral Ray# | 0.11 | 35 | 0.5263 |
| Pectoral Area | 0.58 | 35 | 0.0001 |
| Pectoral Base Length | 0.18 | 35 | 0.2978 |
| Mean Pectoral Fin Ray Length | 0.73 | 35 | 2.22E-07 |
| Pectoral Ray 1 Length | 0.25 | 35 | 0.1306 |
| Pectoral Ray 2 Length | 0.46 | 35 | 0.0044 |
| Pectoral Ray 3 Length | 0.54 | 35 | 0.0006 |
| Pectoral Ray 4 Length | 0.55 | 35 | 0.0004 |
| Pectoral Ray 5 Length | 0.65 | 35 | 1.40E-05 |
| Pectoral Ray 6 Length | 0.70 | 35 | 1.28E-06 |
| Pectoral Ray 7 Length | 0.62 | 34 | 5.04E-05 |
| Pectoral Ray 8 Length | 0.52 | 33 | 0.0012 |
| Pectoral Ray 9 Length | 0.52 | 31 | 0.0017 |
| Pectoral Ray 10 Length | 0.42 | 22 | 0.0434 |
| Pectoral Ray 11 Length | 0.56 | 11 | 0.0450 |
| Pectoral Ray 12 Length | 0.86 | 3 | 0.0648 |

The overarching goal of this study was to determine which fins had the highest levels of sexual dimorphism. Anal and dorsal fins were much more sexually dimorphic than caudal, pelvic, or pectoral fins (Tables 3-4, Figures 3-5). On average, dorsal and anal fin areas were 68% and 55% larger, respectively, in males than in females. Average dorsal and anal fin ray lengths were particularly dimorphic (3.84 and 3.94, respectively). On average, anal and dorsal fin traits were 3X more dimorphic than caudal, pectoral, or pelvic fins. Figure 4 shows the raw and size-adjusted values for the dorsal and anal fin traits. Fin area, fin ray length, and fin base length all showed signs of high sexual dimorphism, suggesting that sexual dimorphism in both base length and ray length contributes to the strong signature of sexual dimorphism in fin area.

**Figure 3.**
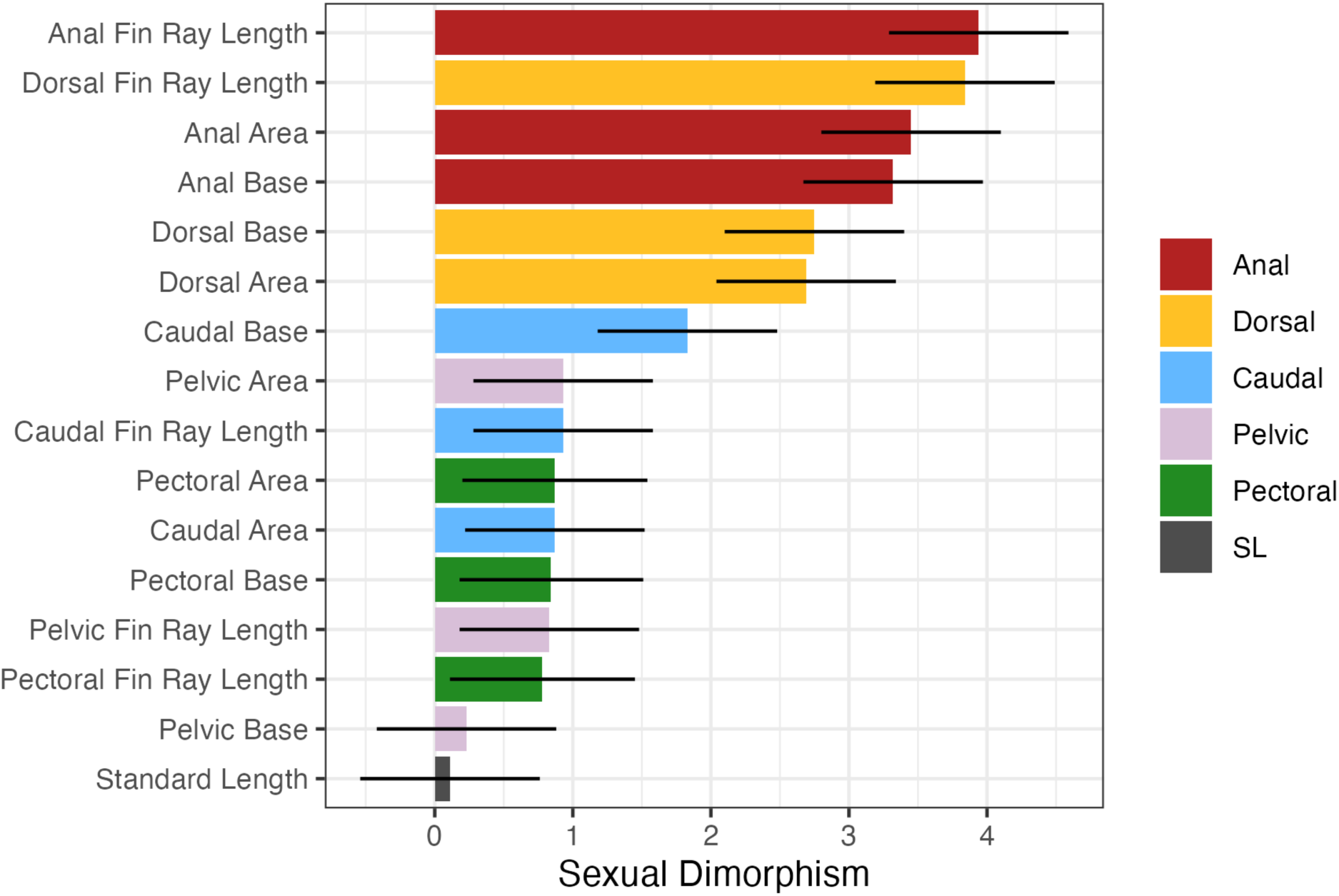
Sexual dimorphism for anal, dorsal, caudal, pelvic, and pectoral fins and standard length. Effect sizes and 95% confidence intervals are shown.

**Figure 4.**
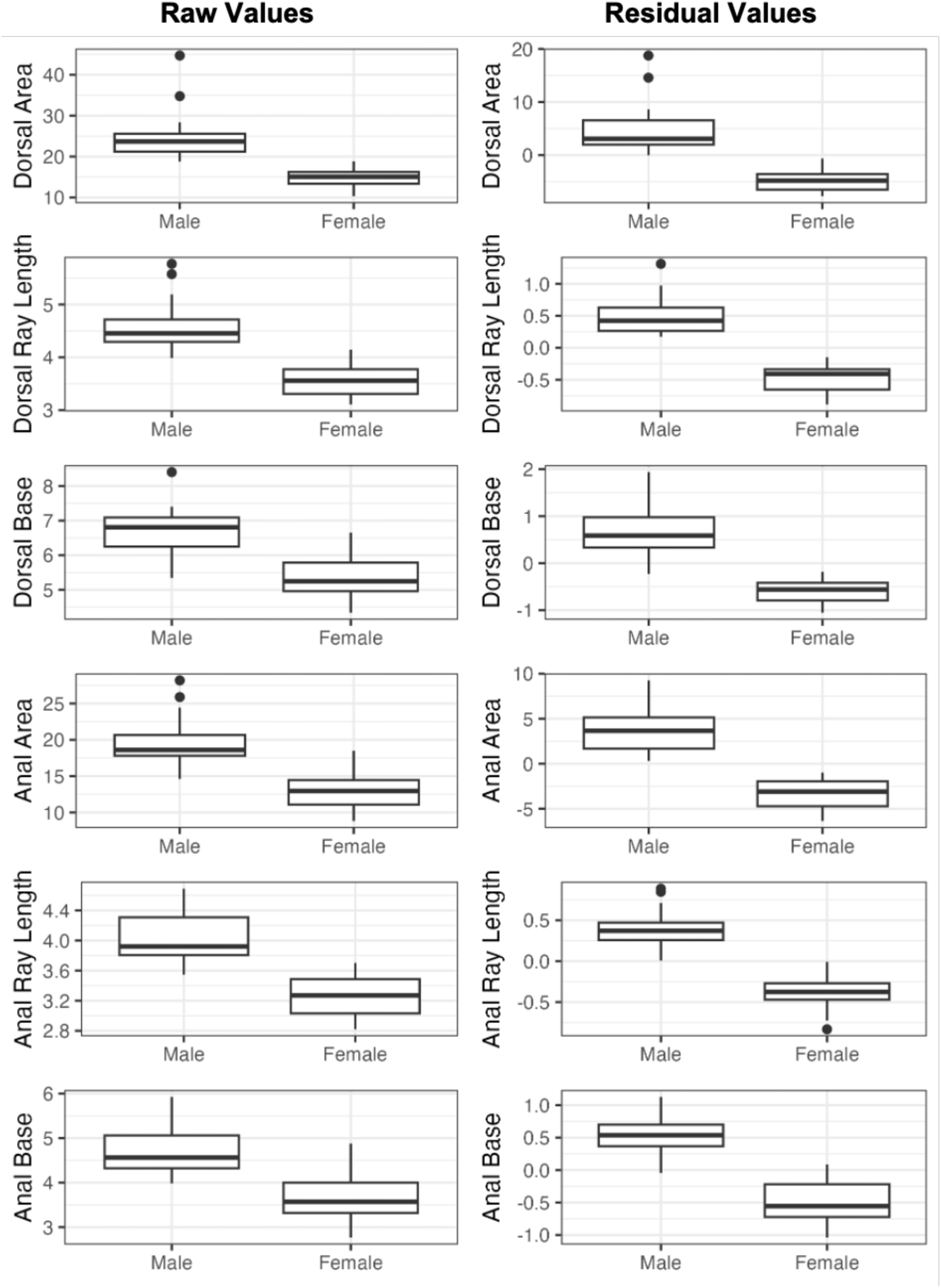
Raw and residual values for dorsal and anal fin traits as a function of sex. Residual values are from a linear regression of raw trait values on standard length.

**Figure 5.**
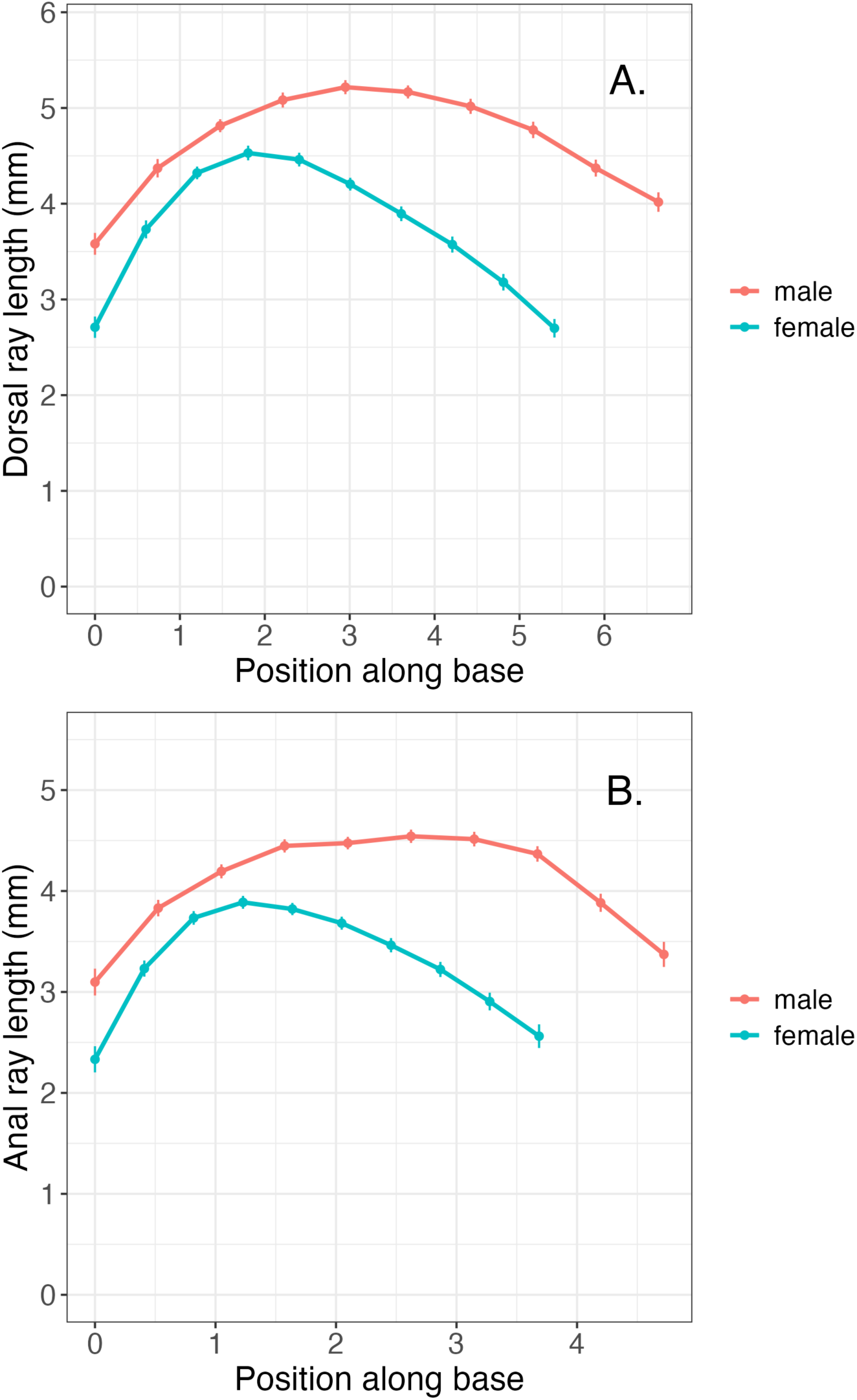
Least-square means ± SE of ray lengths versus relative position along the fin base relative to the insertion point (0) and the sex-specific average fin base length.

**Table 3.** Analyses of variance as a function of sex and population for size corrected traits. All trait values are residuals from a linear regression on standard length. F-values are listed on top and p-values below. Values in bold and italic remain statistically significant after a sequential bonferonni (’holm’) adjustment. The levels of sexual dimorphism are calculated as the effect size due to sex plus their 95% confidence limits.

| Trait | Model | Sex | Population | Sex Dimorph (95% CL) |
| --- | --- | --- | --- | --- |
| Resid Dorsal Area | **35.27**<br>**3.30E-09** | **70.42**<br>**5.39E-10** | 0.01<br>0.9083 | 2.69 (2.04,3.34) |
| Resid Dorsal Base | **36.99**<br>**1.86E-09** | **73.74**<br>**3.08E-10** | 0.07<br>0.7862 | 2.75 (2.1,3.4) |
| Resid Dorsal Fin Ray Length | **71.98**<br>**2.63E-13** | **143.58**<br>**4.01E-14** | 0.87<br>0.3583 | 3.84 (3.19,4.49) |
| Resid Anal Base | **53.73**<br>**1.56E-11** | **107.25**<br>**2.42E-12** | 0.03<br>0.8605 | 3.32 (2.67,3.97) |
| Resid Anal Area | **57.92**<br>**5.60E-12** | **115.74**<br>**8.51E-13** | 0.00<br>0.9721 | 3.45 (2.8,4.1) |
| Resid Anal Fin Ray Length | **76.81**<br>**1.03E-13** | **150.86**<br>**1.95E-14** | 3.95<br>0.0546 | 3.94 (3.29,4.59) |
| Resid Caudal Base | **16.67**<br>**7.51E-06** | **32.44**<br>**1.77E-06** | 0.63<br>0.4309 | 1.83 (1.18,2.48) |
| Resid Caudal Area | 3.88<br>0.0297 | 7.4<br>0.0100 | 0.29<br>0.5966 | 0.87 (0.22,1.52) |
| Resid Caudal Fin Ray Length | 4.54<br>0.0174 | **8.46**<br>**0.0062** | 0.75<br>0.3910 | 0.93 (0.28,1.58) |
| Resid Pelvic Base | 0.73<br>0.4906 | 0.53<br>0.4717 | 0.96<br>0.3337 | 0.23 (-0.42,0.88) |
| Resid Pelvic Area | 4.32<br>0.0208 | **8.42**<br>**0.0063** | 0.15<br>0.6993 | 0.93 (0.28,1.58) |
| Resid Pelvic Fin Ray Length | 4.22<br>0.0226 | 6.68<br>0.0139 | 1.94<br>0.1720 | 0.83 (0.18,1.48) |
| Pectoral Base | 3.44<br>0.0437 | 6.59<br>0.0148 | 0.37<br>0.5459 | 0.84 (0.18,1.51) |
| Pectoral Area | 3.58<br>0.0388 | 7.03<br>0.0121 | 0.19<br>0.6658 | 0.87 (0.2,1.54) |
| Pectoral Fin Ray Length | 3.14<br>0.0558 | 5.62<br>0.0235 | 0.56<br>0.4610 | 0.78 (0.11,1.45) |

Denominator DF are 36 for all models except for pectoral fin traits where they are 34.

**Table 4.** Analyses of variance on individual size-adjusted fin ray lengths. Adjusted-fin ray lengths were residuals from a regression on standard length. F-values in **bold** remain statistically significant after a sequential bonferonni (’holm’) correction. Values in plain italics

| **Trait** | **Model F** | **Sex F** | **Source F** | **Sex Dimorphism** | **Denom DF** |
| --- | --- | --- | --- | --- | --- |
| Resid Dorsal Ray 1 | **14.89** | **29.76** | 0.00 | 1.75 (1.1, 2.4) | 36 |
| Resid Dorsal Ray 2 | **11.33** | **22.63** | 0.10 | 1.52 (0.87, 2.17) | 36 |
| Resid Dorsal Ray 3 | **13.44** | **26.89** | 0.01 | 1.66 (1.01, 2.31) | 36 |
| Resid Dorsal Ray 4 | **12.78** | **25.55** | 0.06 | 1.62 (0.97, 2.27) | 36 |
| Resid Dorsal Ray 5 | **27.33** | **54.6** | 0.17 | 2.37 (1.72, 3.02) | 36 |
| Resid Dorsal Ray 6 | **51.76** | **103.4** | 0.39 | 3.26 (2.61, 3.91) | 36 |
| Resid Dorsal Ray 7 | **52.72** | **105.24** | 0.53 | 3.29 (2.64, 3.94) | 36 |
| Resid Dorsal Ray 8 | **50.35** | **100.58** | 0.38 | 3.21 (2.56, 3.86) | 36 |
| Resid Dorsal Ray 9 | **47.3** | **94.37** | 0.55 | 3.11 (2.46, 3.76) | 36 |
| Resid Dorsal Ray 10 | **47.47** | **94.21** | 1.95 | 3.16 (2.5, 3.82) | 35 |
| Resid Anal Ray 1 | **8.87** | **17.36** | 0.52 | 1.34 (0.69, 1.99) | 36 |
| Resid Anal Ray 2 | **14.29** | **28.12** | 0.67 | 1.7 (1.05, 2.35) | 36 |
| Resid Anal Ray 3 | **12.09** | **23.51** | 0.89 | 1.55 (0.9, 2.2) | 36 |
| Resid Anal Ray 4 | **21.48** | **40.69** | 2.80 | 2.04 (1.39, 2.69) | 36 |
| Resid Anal Ray 5 | **30.17** | **59.31** | 1.49 | 2.47 (1.82, 3.12) | 36 |
| Resid Anal Ray 6 | **44.74** | **89.05** | 0.81 | 3.02 (2.37, 3.67) | 36 |
| Resid Anal Ray 7 | **54.35** | **108.63** | 0.29 | 3.34 (2.69, 3.99) | 36 |
| Resid Anal Ray 8 | **58.65** | **116.85** | 0.91 | 3.46 (2.81, 4.11) | 36 |
| Resid Anal Ray 9 | **30.81** | **61.44** | 0.41 | 2.51 (1.86, 3.16) | 36 |
| Resid Anal Ray 10 | **13.64** | **26.19** | 2.81 | 1.77 (1.06, 2.48) | 31 |
| Resid Median Caudal Ray | 1.86 | 3.01 | 0.80 | 0.56 (-0.09, 1.21) | 36 |
| Resid Caudal Dorsal Ray 1 | 2.11 | 3.51 | 0.79 | 0.6 (-0.05, 1.25) | 36 |
| Resid Caudal Dorsal Ray 2 | 1.18 | 1.9 | 0.51 | 0.44 (-0.21, 1.09) | 36 |
| Resid Caudal Dorsal Ray 3 | 2.22 | 3.95 | 0.58 | 0.64 (-0.01, 1.29) | 36 |
| Resid Caudal Dorsal Ray 4 | 1.27 | 2.46 | 0.10 | 0.5 (-0.15, 1.15) | 36 |
| Resid Caudal Dorsal Ray 5 | 1.48 | 2.43 | 0.59 | 0.5 (-0.15, 1.15) | 36 |
| Resid Caudal Dorsal Ray 6 | 0.82 | 1.15 | 0.53 | 0.34 (-0.31, 0.99) | 36 |
| Resid Caudal Ventral Ray 1 | 2.51 | 4.82 | 0.25 | 0.7 (0.05, 1.35) | 36 |
| Resid Caudal Ventral Ray 2 | 1.24 | 2.29 | 0.23 | 0.49 (-0.16, 1.14) | 36 |
| Resid Caudal Ventral Ray 3 | 1.4 | 2.53 | 0.31 | 0.51 (-0.14, 1.16) | 36 |
| Resid Caudal Ventral Ray 4 | 2.32 | **4.44** | 0.25 | 0.68 (0.03, 1.33) | 36 |
| Resid Caudal Ventral Ray 5 | 6.23 | **12.15** | 0.43 | **1.12 (0.47, 1.77)** | 36 |
| Resid Caudal Ventral Ray 6 | **6.74** | **12.22** | 1.47 | **1.12 (0.47, 1.77)** | 36 |
| Resid Pelvic Ray 1 | 2.93 | 1.92 | 4.09 | 0.44 (-0.21, 1.09) | 36 |
| Resid Pelvic Ray 2 | 2.41 | 2.19 | 2.76 | 0.47 (-0.18, 1.12) | 36 |
| Resid Pelvic Ray 3 | **8.26** | **15.65** | 1.08 | **1.27 (0.62, 1.92)** | 36 |
| Resid Pelvic Ray 4 | 5.17 | 9.8 | 0.67 | 1 (0.35, 1.65) | 36 |
| Resid Pelvic Ray 5 | 2.16 | 4.3 | 0.03 | 0.66 (0.01, 1.31) | 36 |
| Resid Pelvic Ray 6 | 0.41 | 0.6 | 0.26 | 0.27 (-0.43, 0.97) | 31 |
| Resid Pectoral Ray 1 | 0.03 | 0 | 0.07 | -0.02 (-0.69, 0.65) | 34 |
| Resid Pectoral Ray 2 | 0.89 | 0.82 | 0.91 | 0.3 (-0.37, 0.97) | 34 |
| Resid Pectoral Ray 3 | 2.74 | 4.13 | 1.20 | 0.67 (0, 1.34) | 34 |
| Resid Pectoral Ray 4 | 2.19 | 3.75 | 0.54 | 0.64 (-0.03, 1.31) | 34 |
| Resid Pectoral Ray 5 | **7.93** | **14.85** | 0.80 | **1.27 (0.6, 1.94)** | 34 |
| Resid Pectoral Ray 6 | 4.19 | 7.96 | 0.31 | 0.93 (0.26, 1.6) | 34 |
| Resid Pectoral Ray 7 | 0.9 | 1.69 | 0.11 | 0.43 (-0.25, 1.11) | 33 |
| Resid Pectoral Ray 8 | 0.91 | 1.8 | 0.02 | 0.45 (-0.23, 1.14) | 32 |
| Resid Pectoral Ray 9 | 1.18 | 2.33 | 0.00 | 0.53 (-0.18, 1.25) | 30 |

Anal and dorsal fins were particularly dimorphic for the posterior fin rays. Table 4 shows the levels of sexual dimorphism for each fin ray. Posterior dorsal and anal fin ray lengths had sexual dimorphism levels that were ∼2X higher than those for some anterior fin rays. Figure 5 shows the least square means for each fin ray versus the relative position of each ray given the fin base length as a function of sex. The enlarged posterior dorsal and anal fin rays positioned further along the body axis make the fin area larger, particularly on the posterior portion of the fin.

Caudal, pelvic, and pectoral fins also showed moderate levels of sexual dimorphism, with males having larger fin areas and longer fin rays than females. Males also had larger caudal and pectoral fin base lengths than did females. These effects were modest.

This study also asked whether the levels of variability vary as a function of sex, particularly for traits with high levels of sexual dimorphism. Overall, the levels of variability were remarkably similar between males and females across traits (Table 5, Figure 4). Coefficients of variation ranged from ∼0.05 for average fin ray lengths to ∼0.20 for some fin areas. The levels of variability were not consistently higher for traits with high levels of sexual dimorphism.

**Table 5.** Means, Coefficients of variation on raw values, and Coefficients of Variation on Size-Corrected Values for males and females.

| **Trait** | **Mean**<br>**(Male)** | **Mean**<br>**(Female)** | **CV**<br>**(Male)** | **CV**<br>**(Female)** | **Size-**<br>**Corrected**<br>**CV**<br>**(Male)** | **Size-**<br>**Corrected**<br>**CV**<br>**(Female)** |
| --- | --- | --- | --- | --- | --- | --- |
| ***Ray Numbers*** |  |  |  |  |  |  |
| Dorsal | 10.49 | 10.84 | 0.069 | 0.064 | 0.067 | 0.055 |
| Anal | 10.05 | 10.38 | 0.073 | 0.093 | 0.072 | 0.085 |
| Caudal | 13.47 | 13.80 | 0.045 | 0.050 | 0.045 | 0.049 |
| Pelvic | 5.78 | 5.85 | 0.061 | 0.079 | 0.060 | 0.078 |
| Pectoral | 9.79 | 9.42 | 0.134 | 0.134 | 0.134 | 0.132 |
| ***Fin Area*** |  |  |  |  |  |  |
| Dorsal | 24.71 | 14.69 | 0.249 | 0.159 | 0.190 | 0.100 |
| Anal | 19.74 | 12.72 | 0.173 | 0.205 | 0.120 | 0.118 |
| Caudal | 44.87 | 38.97 | 0.153 | 0.258 | 0.133 | 0.154 |
| Pelvic | 7.63 | 6.09 | 0.237 | 0.313 | 0.156 | 0.297 |
| Pectoral | 17.57 | 14.62 | 0.263 | 0.246 | 0.206 | 0.191 |
| ***Base length*** |  |  |  |  |  |  |
| Dorsal | 6.65 | 5.40 | 0.111 | 0.123 | 0.085 | 0.047 |
| Anal | 4.73 | 3.67 | 0.120 | 0.153 | 0.065 | 0.083 |
| Caudal | 4.40 | 4.05 | 0.097 | 0.103 | 0.038 | 0.046 |
| Pelvic | 1.32 | 1.27 | 0.178 | 0.204 | 0.162 | 0.197 |
| Pectoral | 2.02 | 1.77 | 0.172 | 0.131 | 0.167 | 0.130 |
| ***Mean Fin Ray***<br>***Length*** |  |  |  |  |  |  |
| Dorsal | 4.58 | 3.59 | 0.106 | 0.090 | 0.063 | 0.052 |
| Anal | 4.05 | 3.25 | 0.093 | 0.081 | 0.054 | 0.055 |
| Caudal | 5.51 | 5.25 | 0.086 | 0.100 | 0.041 | 0.053 |
| Pelvic | 3.20 | 2.87 | 0.107 | 0.193 | 0.060 | 0.181 |
| Pectoral | 3.83 | 3.57 | 0.138 | 0.118 | 0.095 | 0.068 |
| ***Dorsal Fin Ray***<br>***Lengths*** |  |  |  |  |  |  |
| Ray 1 | 3.59 | 2.71 | 0.179 | 0.148 | 0.166 | 0.134 |
| Ray 2 | 4.38 | 3.73 | 0.131 | 0.121 | 0.105 | 0.097 |
| Ray 3 | 4.82 | 4.32 | 0.085 | 0.093 | 0.053 | 0.074 |
| Ray 4 | 5.09 | 4.52 | 0.096 | 0.091 | 0.072 | 0.067 |
| Ray 5 | 5.23 | 4.45 | 0.105 | 0.089 | 0.067 | 0.059 |
| Ray 6 | 5.18 | 4.20 | 0.100 | 0.097 | 0.059 | 0.063 |
| Ray 7 | 5.03 | 3.89 | 0.112 | 0.113 | 0.076 | 0.073 |
| Ray 8 | 4.78 | 3.57 | 0.117 | 0.132 | 0.086 | 0.091 |
| Ray 9 | 4.38 | 3.17 | 0.139 | 0.159 | 0.094 | 0.108 |
| Ray 10 | 4.02 | 2.70 | 0.181 | 0.226 | 0.107 | 0.152 |
| Ray 11 | 3.36 | 2.27 | 0.221 | 0.217 | 0.077 | 0.157 |
| ***Anal Fin Ray Lengths*** |  |  |  |  |  |  |
| Ray 1 | 3.10 | 2.33 | 0.181 | 0.257 | 0.169 | 0.257 |
| Ray 2 | 3.83 | 3.23 | 0.113 | 0.108 | 0.078 | 0.107 |
| Ray 3 | 4.20 | 3.73 | 0.089 | 0.085 | 0.064 | 0.081 |
| Ray 4 | 4.45 | 3.88 | 0.084 | 0.084 | 0.055 | 0.076 |
| Ray 5 | 4.48 | 3.82 | 0.086 | 0.079 | 0.063 | 0.064 |
| Ray 6 | 4.55 | 3.68 | 0.091 | 0.090 | 0.068 | 0.069 |
| Ray 7 | 4.52 | 3.46 | 0.101 | 0.114 | 0.076 | 0.080 |
| Ray 8 | 4.37 | 3.22 | 0.119 | 0.132 | 0.077 | 0.097 |
| Ray 9 | 3.89 | 2.90 | 0.132 | 0.168 | 0.105 | 0.126 |
| Ray 10 | 3.36 | 2.57 | 0.183 | 0.222 | 0.168 | 0.157 |
| Ray 11 | 2.96 | 2.65 | 0.146 | 0.165 | 0.129 | 0.117 |
| ***Caudal Ray Lengths*** |  |  |  |  |  |  |
| Dorsal Ray 7 | 4.40 | 4.40 | 0.147 | 0.156 | 0.096 | 0.080 |
| Dorsal Ray 6 | 5.19 | 5.01 | 0.112 | 0.150 | 0.079 | 0.103 |
| Dorsal Ray 5 | 5.64 | 5.44 | 0.097 | 0.126 | 0.050 | 0.077 |
| Dorsal Ray 4 | 5.76 | 5.55 | 0.096 | 0.118 | 0.056 | 0.075 |
| Dorsal Ray 3 | 5.72 | 5.52 | 0.089 | 0.098 | 0.045 | 0.057 |
| Dorsal Ray 2 | 5.63 | 5.50 | 0.086 | 0.092 | 0.044 | 0.050 |
| Dorsal Ray 1 | 5.58 | 5.39 | 0.089 | 0.097 | 0.048 | 0.061 |
| Median Ray | 5.60 | 5.44 | 0.083 | 0.093 | 0.043 | 0.050 |
| Ventral Ray 1 | 5.62 | 5.39 | 0.075 | 0.109 | 0.040 | 0.066 |
| Ventral Ray 2 | 5.62 | 5.48 | 0.076 | 0.093 | 0.046 | 0.043 |
| Ventral Ray 3 | 5.65 | 5.50 | 0.082 | 0.092 | 0.046 | 0.046 |
| Ventral Ray 4 | 5.67 | 5.47 | 0.083 | 0.091 | 0.049 | 0.050 |
| Ventral Ray 5 | 5.49 | 5.11 | 0.095 | 0.102 | 0.055 | 0.066 |
| Ventral Ray 6 | 4.91 | 4.36 | 0.108 | 0.168 | 0.074 | 0.132 |
| ***Pelvic Ray Lengths*** |  |  |  |  |  |  |
| Ray 1 | 2.61 | 2.41 | 0.136 | 0.248 | 0.109 | 0.240 |
| Ray 2 | 3.30 | 3.09 | 0.126 | 0.195 | 0.079 | 0.182 |
| Ray 3 | 3.88 | 3.37 | 0.087 | 0.167 | 0.055 | 0.152 |
| Ray 4 | 3.78 | 3.21 | 0.140 | 0.219 | 0.090 | 0.212 |
| Ray 5 | 3.12 | 2.73 | 0.173 | 0.248 | 0.137 | 0.241 |
| Ray 6 | 2.44 | 2.29 | 0.191 | 0.292 | 0.184 | 0.280 |
| ***Pectoral Ray***<br>***Lengths*** |  |  |  |  |  |  |
| Ray 1 | 2.51 | 2.51 | 0.390 | 0.350 | 0.382 | 0.333 |
| Ray 2 | 3.63 | 3.42 | 0.200 | 0.218 | 0.173 | 0.197 |
| Ray 3 | 4.33 | 3.95 | 0.157 | 0.162 | 0.136 | 0.127 |
| Ray 4 | 4.67 | 4.34 | 0.117 | 0.148 | 0.106 | 0.109 |
| Ray 5 | 4.91 | 4.41 | 0.097 | 0.140 | 0.069 | 0.097 |
| Ray 6 | 4.73 | 4.34 | 0.119 | 0.143 | 0.086 | 0.091 |
| Ray 7 | 4.35 | 4.09 | 0.194 | 0.161 | 0.161 | 0.107 |
| Ray 8 | 3.81 | 3.49 | 0.236 | 0.222 | 0.205 | 0.179 |
| Ray 9 | 3.18 | 2.80 | 0.346 | 0.291 | 0.277 | 0.249 |
| Ray 10 | 2.48 | 2.49 | 0.399 | 0.310 | 0.347 | 0.291 |

Size-corrected anal fin and dorsal fin traits were strongly correlated with one another in both sexes. Size-adjusted dorsal and anal fin area was strongly correlated in males (Table 6, Figure 6, R_17_ = 0.84, 95% CL: 0.94, 0.62) and moderately correlated in females (Table 6). There were positive correlations among most of the fin areas for both sexes, but only the correlation between size-corrected dorsal and anal fin area remained statistically significant after a sequential Bonferroni correction in males. Furthermore, the 95% confidence intervals for the correlation between dorsal and anal fins excluded the other correlation coefficients between other fin area traits. The correlation between dorsal and anal fin area was also high in females (and was the highest correlation among fin areas), but it was not statistically significant after a sequential Bonferroni correction (Table 6, Figure 6).

**Figure 6.**
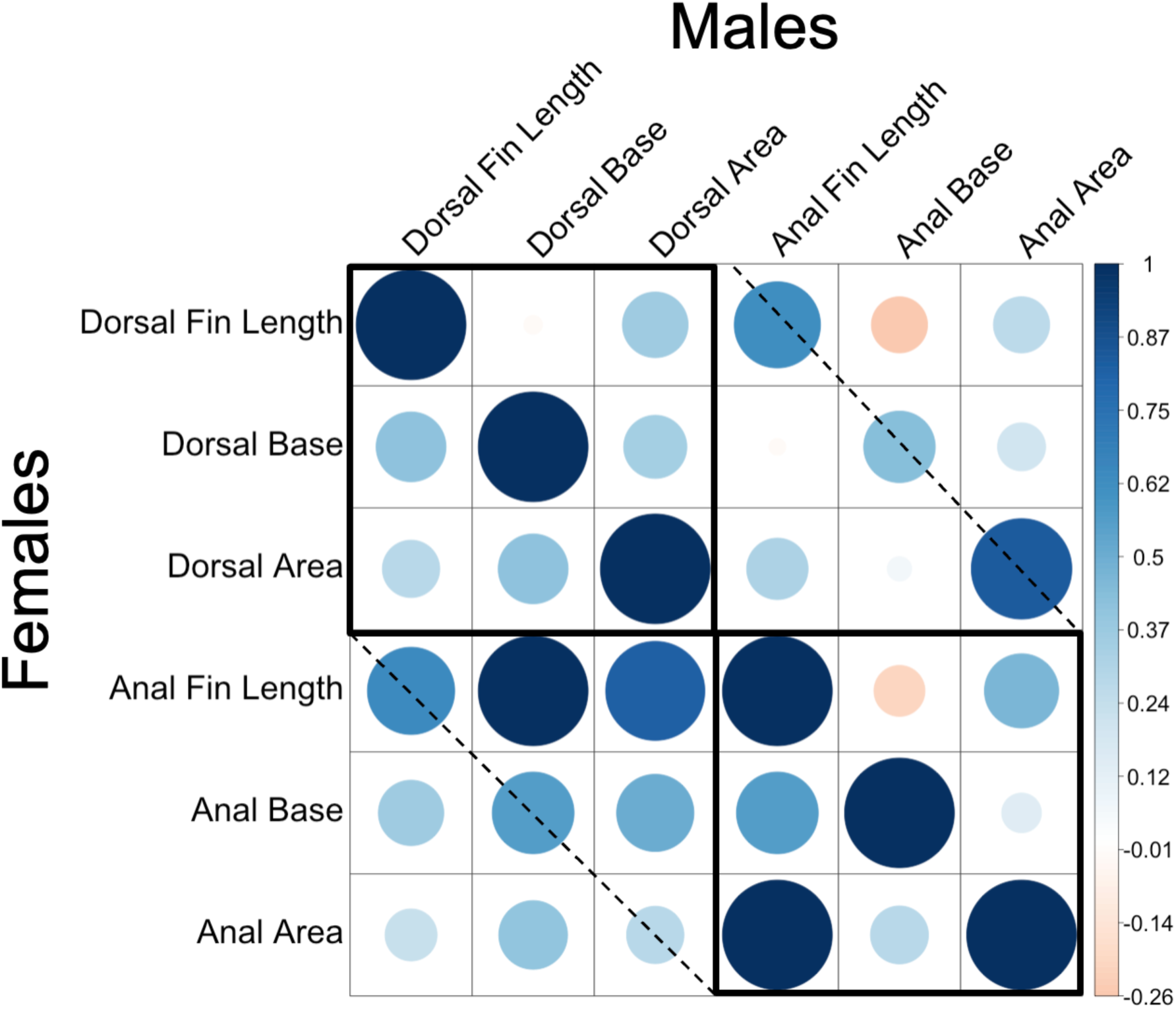
Correlations between fin ray length, fin base length, and fin area in dorsal and anal fins for males (top) and females bottom). Bold squares show correlations between elements within a fin type. The dashed line shows the correlation between the same fin trait (i.e., fin ray lengths) between the dorsal and anal fins.

**Table 6.** Pearson correlation coefficients between size-corrected dorsal, anal, caudal, pelvic, and pectoral fin traits (A: Fin Area, B: Average Fin Length, C: Base Length). Values above the diagonal are correlations across males. Values below the diagnonal are correlations across females.

|  | Dorsal Area | Anal Area | Caudal Area | Pelvic Area | Pectoral Area |
| --- | --- | --- | --- | --- | --- |
| Dorsal Area |  | **0.84** | 0.59 | 0.49 | 0.37 |
| Anal Area | 0.51 |  | 0.51 | 0.47 | 0.34 |
| Caudal Area | 0.16 | 0.33 |  | 0.42 | 0.04 |
| Pectoral Area | 0.22 | 0.54 | -0.04 |  | 0.51 |
| Pelvic Area | 0.12 | 0.28 | 0.2 | 0.29 |  |

B. Average Fin Length
|  | Dorsal Fin L | Anal Fin L | Caudal Fin L | Pelvic Fin L | Pectoral Fin L |
| --- | --- | --- | --- | --- | --- |
| Dorsal Fin L |  | **0.62** | 0.29 | 0.43 | 0.04 |
| Anal Fin L | **0.63** |  | 0.37 | 0.4 | 0.21 |
| Caudal Fin L | 0.47 | 0.45 |  | 0.06 | 0.05 |
| Pelvic Fin L | 0.47 | 0.59 | 0.35 |  | 0.06 |
| Pectoral Fin L | 0.41 | 0.01 | 0.1 | 0.41 |  |

C. Base Length Correlations
|  | Dorsal Base | Anal Base | Caudal Base | Pelvic Base | Pectoral Base |
| --- | --- | --- | --- | --- | --- |
| Dorsal Base |  | 0.43 | 0.48 | 0.1 | -0.07 |
| Anal Base | **0.82** |  | 0.15 | 0 | -0.02 |
| Caudal Base | -0.1 | -0.33 |  | -0.07 | 0.15 |
| Pelvic Base | 0.34 | 0.45 | -0.15 |  | -0.34 |
| Pectoral Base | 0.34 | 0.22 | 0.09 | -0.08 |  |

Similar patterns were seen for fin ray lengths and base lengths (Table 6, Figure 6). Dorsal and anal fin ray lengths were strongly correlated with one another in both males and females, and both remained statistically significant after a sequential Bonferroni correction. For both males and females, the correlations between dorsal and anal fin ray lengths were higher than the correlations among fin ray lengths for other combinations of fins. Females also had a high correlation between dorsal and anal fin base lengths (R_18_ = 0.84), which was nearly 2X higher than any of the other correlations for base lengths. Among males, there were positive correlations between dorsal and caudal fin base lengths, but they did not remain statistically significant after a sequential Bonferroni correction. While dorsal fin traits (i.e., fin ray length, fin area) were correlated with anal fin traits, there were not strong correlations between fin ray length and fin base length, particularly in males (Figure 6).

## DISCUSSION

In this study, we sought to determine the degree of sexual dimorphism in fin morphology in the bluefin killifish, *Lucania goodei.* The results paint a picture of high sexual dimorphism in dorsal and anal fins relative to the other fins (pelvic, pectoral, and caudal fins). The dimorphism was particularly pronounced in the posterior fin rays for both the dorsal and anal fins. The result is in keeping with the findings of Davis et al (2025). Davis et al. (2025) measured sexual dimorphism in dorsal, anal, and caudal fins across 20 species of Fundulidae, including bluefin killifish, and found high levels of dimorphism in dorsal and anal fins and lower levels in caudal fins. While the concordance is not surprising, it is reassuring. The present study builds on prior work by using fresh specimens, a larger sample size, and by incorporating measurements of the paired fins (i.e., pectoral and pelvic fins). The larger sample size also enabled investigation of trait correlations within each sex.

One of the major patterns to emerge is that dorsal and anal fins have much higher levels of sexual dimorphism than those found in caudal, pectoral, and pelvic fins. Similar results have been found in other groups. Mainero et al. (2023) performed a similar study in the Arabian killifish, *Aphaniops stoliczkanus,* from the family Aphaniidae, and found a similar pattern with high sexual dimorphism in dorsal and anal fins and lower levels in pectoral and pelvic fins. In a beach-spawning capelin, *Mallotus villosus,* sexual dimorphism was found to be particularly high in the anal fins, but other fins were less so (Orbach, et al., 2019). Englmaier et al. (2022) found apparent high levels of sexual dimorphism in dorsal and anal fins and lower levels in other fins. Elevated dimorphism in dorsal and anal fins has been documented across a range of lineages, including gars, blennies, minnows, cichlids, and salmonids (Chervinski, 1965; McCart, 1965; Ostrand, et al., 2001; McGrath and Hilton, 2012; Englmaier, et al., 2022). This strikingly consistent pattern suggests that several distantly related species face similar selection pressures.

The question of why dorsal and anal fins frequently show such high levels of sexual dimorphism is not fully resolved. Dorsal and anal fins are often involved in signaling between males and females (Thompson and Sturm, 1965; Fuller, 2001; McGhee, et al., 2007; Johnson and Fuller, 2015; Zhou, et al., 2015; Mitchem, et al., 2018). However, a more compelling answer may be that many of these species pair spawn. In many species, males use their dorsal and anal fins to clasp females and guide gamete release (Newman, 1907; Foster, 1967; Arndt, 1971; Able and Hata, 1984). The anal fin, in particular, may play a large role in controlling the direction of the flow of sperm and eggs, and preventing sperm of other males from reaching the eggs (Barbas and Gilg, 2018). Experimental work supports this idea: in medaka, shortening the anal fin reduces both fertilization success and male attractiveness to females (Koseki, et al., 2000; Fujimoto, et al., 2014). Similar results have been found in bluefin killifish, where altering the male anal fin has strong effects on fertilization success and moderate effects on male/male competition and female mating preferences (Smelko and Fuller, in preparation).

More challenging is the question of why the caudal, pectoral, and pelvic fins also exhibit moderate dimorphism in some species. In some taxa, paired fins have clearly evolved mating-related functions. In sticklebacks, for instance, sexual dimorphism in pectoral fins are associated with male parental care (Bakker and Mundwiler, 1999), and experimental reductions in pectoral fin size alter fanning behavior and possibly other life-history traits (Künzler and Bakker, 2000). Pelvic fins are often dramatically dimorphic. In the cichlid *Cyathopharynx fucifer*, males use elongated pelvic fins in courtship displays (Karino, 1997). In one species of ricefish, females brood embryos below their abdomen and have evolved modified pelvic fins to protect the offspring (Flury, et al., 2022). However, in many other species, these fins lack an obvious reproductive function, and the pattern of strong dimorphism in dorsal and anal fins, with weaker or absent dimorphism in other fins, persists.

Why do fins differ in their degree of sexual dimorphism? One possibility is that overall fin growth is weakly correlated across fin types, such that strong selection on dorsal or anal fins produces indirect changes in others. Alternatively, fins may vary in the relative intensity of natural versus sexual selection, leading to differing dimorphism levels. Resolving these possibilities will require direct measurements of selection acting on fin morphology.

This study also found strong correlations between dorsal and anal fin traits in both males and females, suggesting a combination of pleiotropy, shared developmental pathways, and/or correlated selection. Comparative studies have explored the modularity of fin traits, asking whether certain fins evolve in concert while others evolve more independently. Several authors have identified a distinct “dorsal-anal fin module,” in which traits of the dorsal and anal fins are more tightly integrated with one another than with other fins. For example, studies of fin positioning have shown that dorsal and anal fins form a module that is relatively independent from modules involving the pectoral and pelvic fins (Mabee, et al., 2002; Larouche, et al., 2015). The dorsal and anal fin module appears to have been present in Actinopterygians for at least 400 MY (Mabee, et al., 2002).

Further support for shared developmental control comes from QTL studies in medaka. Kawajiri et al. (2014) crossed two medaka populations (one exhibiting low and the other high sexual dimorphism in dorsal and anal fin size) and found that nearly all QTL for dorsal fin traits co-localized with QTL for anal fin traits. Although some additional QTL were unique to the anal fin, the overlap strongly supports pleiotropic effects in medaka. Similar genetic studies have not yet been conducted in bluefin killifish. However, developmental studies show that dorsal and anal fins emerge simultaneously during embryogenesis and follow similar developmental trajectories during early life stages (Crawford and Balon, 1994a, c, b). Further work is needed to determine the extent to which genetic correlations due to pleiotropy occur in bluefin killifish.

A key question, however, is how sexual dimorphism arises within these shared developmental modules. Several lines of evidence suggest that fin growth is sensitive to sex hormones. In medaka, dorsal and anal fins follow similar developmental patterns early in ontogeny but diverge between sexes at the same time that sex hormone levels begin to differ (Kawajiri, et al., 2014; Kawajiri, et al., 2015; Zhang, et al., 2023). Furthermore, androgen treatments in females can induce male-specific traits, such as the papillary process typically found on the anal fins in male medaka (Ogino, et al., 2014). In guppies, mosquitofish, and platyfish, treating females with androgens causes their anal fin to elongate and form a structure that resembles a gonopodium (Turner, 1942; Sangster, 1948; Hopper, 1949; Angus, et al., 2001; Offen, et al., 2013). In bluefin killifish, Fuller and Travis (2004) demonstrated that methyltestosterone administration induces male-like coloration patterns in females. Moreover, the presence of androgen, estrogen, and progesterone receptors in the dorsal and anal fins has been documented in several species (Ngamniyom, et al., 2009; Kawajiri, et al., 2014; Zhang, et al., 2023), including bluefin killifish (Karatgi and Fuller, in preparation). These findings suggest that an ancient developmental pathway is likely present and that alteration of growth patterns with sex-hormones possibly allows for sexual dimorphism in these fins.

Finally, this study allowed us to examine how trait variability differs across fin types and between sexes. We found little evidence that traits with high levels of sexual dimorphism were more or less variable than other traits, nor did we find consistent differences in trait variability between males and females. Over time, expectations regarding variability in male secondary sexual traits have shifted. Early models of sexual selection, including both “good genes” and Fisherian frameworks, predicted that strong selection on male traits would erode genetic variation (Charlesworth, 1987; Falconer, 1996). However, later models incorporating condition-dependent expression suggested greater potential for both genetic and environmental variation in male secondary sexual traits (Rowe and Houle, 1996; Houle and Kondrashov, 2002). There has also been ongoing debate about whether males or females should be more variable overall. Historically, female hormonal cycles were thought to increase phenotypic variability (Zajitschek, et al., 2020). In our data, we found little support for differences in trait variability as a function of sex or degree of sexual dimorphism, though our relatively small sample sizes (∼20 individuals per sex) may limit our power to detect such patterns. We note that if sexual dimorphism in fin size arises from hormone-dependent growth, as hypothesized, then male fin traits may be particularly sensitive to fluctuating hormone levels associated with changes in social status or dominance.

In conclusion, we found strikingly high levels of sexual dimorphism in fin area, fin base length, and fin ray lengths of the dorsal and anal fins. Posterior fin rays were particularly dimorphic in both fins. These traits were also strongly correlated across individuals in both sexes: individuals with larger dorsal fins also had larger anal fins, and males with longer dorsal fin rays tended to have longer anal fin rays. Whether these correlations reflect pleiotropy, shared developmental pathways, or strong correlated selection remains unresolved. We also detected moderate levels of sexual dimorphism in the caudal, pelvic, and pectoral fins. Caudal and pelvic fins may play a role in courtship or male–male competition, given that they possess sexual dichromatic coloration. In contrast, the pectoral fins are neither dichromatic nor clearly involved in mating. One explanation is that selection on some fin traits may produce correlated changes in others. Overall, bluefin killifish provide a powerful model for investigating fundamental questions in organismal biology, particularly regarding (1) how genetic and environmental factors influence trait values and their integration, and (2) how natural and sexual selection shape fin morphology, and whether these selective pressures vary across different fin types.

## Acknowledgments

K. Brockelsby and E. Davis were supported by funds from the School of Integrative Biology and the University of Illinois. All procedures were approved by the Institutional Animal Care and Use Committee at the University of Illinois (protocols #23145 and 22112). We thank A. Aka and R. Karatgi for comments that improved the manuscript.

